# Methionine Triggers Metabolic, Transcriptional, and Epigenetic Reprogramming in Arabidopsis Leaves

**DOI:** 10.1101/2025.11.02.686087

**Authors:** Yonatan Yerushalmy, Michal Dafni, Nasrin Rabach, Yael Hacham, Rachel Amir

## Abstract

Methionine (Met) is a central metabolite in plants, as it serves as a precursor for *S*-adenosylmethionine (SAM), a key methyl donor for epigenetic and metabolic processes. Met is also an essential amino acid that limits the nutritional value of plant-based diets. Understanding how altered Met levels affect the metabolome, transcriptome, and epigenetic regulation of plant leaves remains an open challenge. This study investigates the impact of ectopic Met accumulation in SSE Arabidopsis leaves of transgenic lines expressing a deregulated form of *AtCGS* (*AtD-CGS*) under the seed-specific phaseolin promoter. Unexpected activation of the phaseolin promoter in leaves led to *AtD-CGS* expression and variable Met accumulation among progeny, despite genetic homozygosity. High-Met (HM) plants showed elevated amino acid and sugar levels, enrichment of stress-related transcripts, and suppression of Met biosynthetic genes, while Low-Met (LM) plants showed reduced Met levels and increased non-CG DNA methylation, especially in centromeric and promoter regions. Integrated transcriptome and methylome analyses revealed that high Met levels were associated with the upregulation of stress hormone pathways (abscisic acid, jasmonate, salicylic acid, and ethylene), downregulation of key epigenetic regulators (e.g., MET1, CMTs), and broader transcriptional reprogramming. By contrast, low Met (LM) lines displayed similar expression levels of genes as control plants. Our findings reveal a complex regulatory network whereby Met accumulation reprograms metabolism, gene expression, and DNA methylation patterns. These results suggest feedback between sulfur-carbon metabolism, stress adaptation, and epigenetic control, positioning Met as both a nutrient and a signaling hub in plant physiology.

## Introduction

Methionine (Met) is a fundamental metabolite in plant cells. Beyond its well-known role as the initiating amino acid (AA) in mRNA translation, Met serves as a precursor for *S*-adenosylMet (SAM), the universal methyl-group donor for DNA, RNA, histone, lipid, and secondary metabolite methylation (Roje, 2006; Devi et al., 2023). These methylated products play pivotal roles in cellular regulation, development, and stress responses. In human and animal nutrition, Met is one of the most limiting essential amino acids (AAs) in plants, and its scarcity reduces the nutritional value of many crops (Navik et al., 2021). Consequently, metabolic engineering strategies have aimed to enhance Met levels in seeds and vegetative tissues (Amir, 2010).

Met biosynthesis is tightly controlled by cystathionine γ-synthase (CGS), the first committed enzyme of the pathway (Hacham et al., 2006). A deregulated *Arabidopsis* CGS variant (AtD-CGS), lacking 30 AAs in the N-terminal regulatory domain, is less sensitive to feedback inhibition by Met or SAM (Hacham et al., 2006). Seed-specific expression of *AtD-CGS* in *Arabidopsis thaliana*, tobacco (*Nicotiana tabacum*), and soybean (*Glycine max*) increases seed Met content and as well as primary metabolism, including AAs, sugars, and organic acids, which increases starch and protein (Matityahu et al., 2013; Song et al., 2013; Cohen et al., 2014).

To investigate the basis of this nutritionally valuable phenotype, we further analyzed dry seeds from *Arabidopsis* lines expressing *AtD-CGS* under the control of the seed-specific phaseolin promoter (designated SSE plants) (Cohen et al., 2014). Despite the increased accumulation of Met, starch, and proteins, transcriptomic analysis revealed no substantial upregulation of genes directly associated with their biosynthesis (Cohen et al., 2014). Isotope labeling and transcriptome analyses have revealed that many of the metabolites accumulating in seeds originate in leaves and are transported during seed development (Girija et al., 2023). In line with these observations, senescing SSE leaves were found to contain elevated levels of Met, other AAs, and sugars (Girija et al., 2023). Such metabolite accumulation is commonly associated with abiotic stress responses (Salam et al., 2023). The similarity in metabolic profiles between SSE seeds and leaves suggests that *AtD-CGS* is also expressed in leaves. This is unexpected, as the phaseolin promoter is generally regarded as seed-specific, with transcription typically initiated around 12 days after flowering (DAF) in embryos (Chandrasekharan et al., 2003; Fait et al., 2011). However, previous reports have shown that phaseolin promoter activity can be subject to epigenetic regulation and, under certain conditions, may be ectopically activated also in leaves (Sundaram et al., 2013).

In the present study, we focused on SSE leaves because they appear to be the primary source of metabolites that are transported to and accumulated in seeds, thereby enhancing their nutritional value (Girija et al., 2023), an important agronomic trait. Our specific objectives were to: (i) identify the factors underlying the elevated Met levels in SSE leaves; (ii) elucidate the impact of high Met content in leaves on metabolite accumulation; and (iii) evaluate the effects of contrasting Met levels in SSE leaves on the primary metabolome, transcriptome, and methylome.

## Results

### Evidence for phaseolin promoter activity in SSE leaves

Elevated Met levels were detected in senescing leaves of the SSE line (Girija et al., 2023). This unexpected observation may result from activation of the *AtD-CGS* transgene in senescing leaves. To test this hypothesis, *AtD-CGS* transcript abundance was measured in 14, 21, and 28-day-old pre-flowering leaves. Quantitative real-time PCR (qPCR) revealed a 5.2, 4.1, and 4.3-fold increase, respectively, in *AtCGS* transcript levels compared with plants containing only the phaseolin promoter without the *AtD- CGS* transgene (empty vector, EV), which served as a control (Fig. F1a). Using a validated gas chromatography-mass spectrometry (GC-MS) protocol (Cohen et al., 2014) we detected higher Met levels of 2.7 and 2.1-fold at 14 and 21-day-old leaves. The total AA levels increased by 1.8-fold relative to EV at 28-day-old leaves (Fig. F1a). These findings suggest that the phaseolin promoter is unexpectedly active in SSE leaves, likely contributing to the elevated Met accumulation observed in this tissue. To strengthen the assumption that this promoter is sensitive to high concentrations of Met, the expression level of phaseolin gene, which is naturally fused to the phaseolin promoter, was tested. Phaseolin encodes a major storage protein specifically found in bean (*Phaseolus vulgaris*) seeds (López-Pedrouso et al., 2014). The bean stem of a 30-day-old plant was placed in a tube with 5 mM Met, or water (DDW) as a control for 7 hours. The level of Met and the expression level of phaseolin in the leaves were 51-fold and 7-fold higher, respectively, than control (Fig. 1b). These findings suggest that the phaseolin promoter becomes active in the presence of high Met.

**Figure 1.**
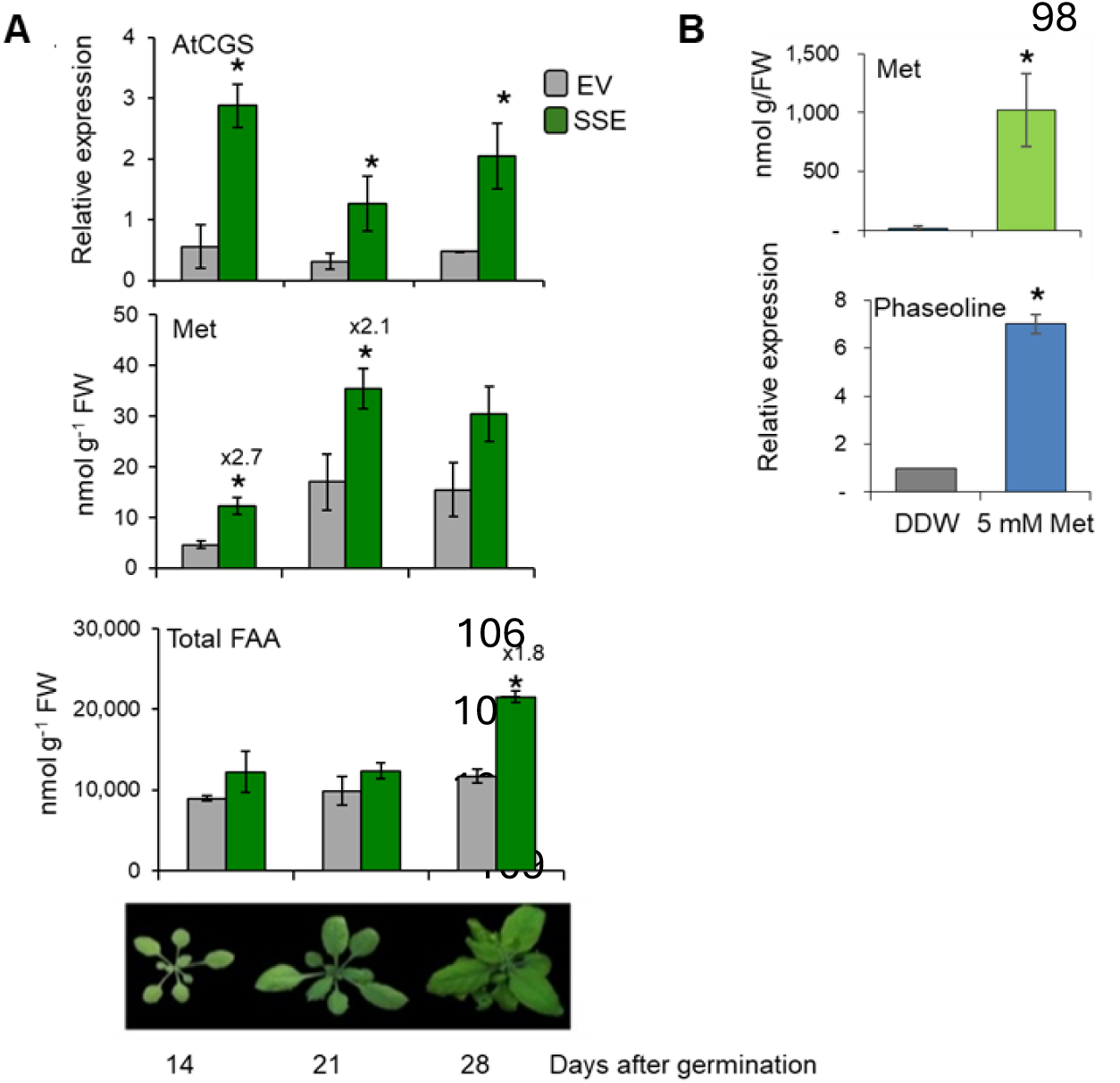
Activation of the seed-specific phaseolin promoter in leaves. (A) Elevated AtCGS expression, Met content, and total free amino acids (T-FAA) in SSE and EV leaves. The expression level of AtCGS was normalized to the endogenous control gene PP2A-A3. Soluble Met levels in leaves (fresh weight [FW]) of EV and SSE plants were quantified via GC-MS and normalized to the ribitol internal standard. Data represent means ± SD from four biological replicates. Statistically significant differences compared to EV (p < 0.05, unpaired Student’s t-test) are indicated by asterisks. (B) Stems of bean plants were supplemented with 5 mM Met or water (DDW) for 7 hours. The expression level phaseolin was normalized to the endogenous control gene Actin. Data represent means ± SD from three biological replicates. Statistically significant differences compared to DDW (p < 0.05, unpaired Student’s t-test) are indicated by asterisks.

### Variation in Met levels among SSE lines of the same generation

In our previous study, we observed that after six generations of SSE, Met levels in seeds declined markedly from approximately sixfold higher than the EV control (Cohen et al., 2014) to only 1.2–1.5- fold higher (Girija et al., 2020). We hypothesize that this reduction may result from progressive epigenetic silencing of regulatory factors controlling Met accumulation in seeds across generations.

To further examine the relationship between Met levels in leaves and seeds of SSE lines, and to better understand the regulatory mechanisms operating in leaves, we generated a new set of transgenic *Arabidopsis* plants expressing the phaseolin::AtD-CGS construct (Cohen et al., 2014). Seeds from 13 independent kanamycin-resistant transformants were screened by immunoblotting (Fig. 1S). Transgenic line No. 35, which exhibited the highest *AtD-CGS* expression (Fig. S2), was selected for further characterization.

Given previous evidence for phaseolin promoter activity in leaves (Fig. 1), we measured leaf Met levels in three T2 progeny lines of line 35. Among them, line 35-3 displayed the highest Met content, with a 2.2-fold increase compared to the EV. In T3 homozygous progeny of line 35-3, Met levels exhibited substantial variability, ranging from 0.2- to 1.9-fold relative to EV (Fig. 2). We selected line 35-3-13, which showed the highest Met accumulation, for self-fertilization. In the T4 generation of this line, leaf Met levels displayed an even wider range, from 0.23- to 2.98-fold relative to EV.

**Figure 2.**
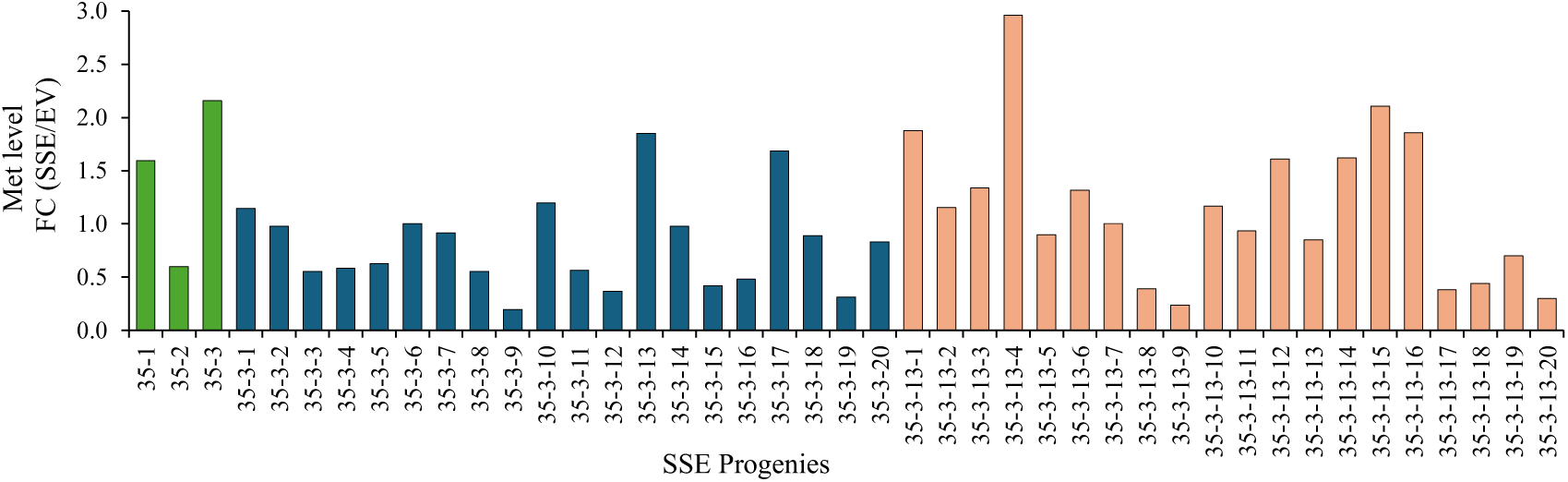
Met levels in leaves of 21-day-old SSE plants across self-fertilized generations T2 (green), T3 (blue), and T4 (pink). The data represents fold changes (FC) in Met content relative to plants harboring the empty vector (EV) control. Met levels were quantified using gas chromatography–mass spectrometry (GC-MS) and normalized to the internal standard ribitol.

### Changes in Met content are associated with alterations in the primary metabolome

Previous studies have shown that, in addition to elevated Met levels, seeds with high Met content often exhibit significant changes in the abundance of other metabolites, particularly soluble AAs and sugars (Matityahu et al., 2013; Song et al., 2013; Cohen et al., 2014; Zhang et al., 2023). To determine whether similar metabolic alterations occur in the leaves of SSE plants, we analyzed T4 homozygous lines exhibiting either high or low Met content. For this analysis, three leaf pools, each comprising material from two individual plants, were prepared to obtain sufficient dry weight for the analyses. The high-Met (HM) pool was generated from lines 35-3-13 no. 1+4, 12+14, and 15+16, whereas the low-Met (LM) pool was generated from lines 35-3-13 no. 8+9, 17+18, and 19+20. Met levels were measured in these pools, revealing that HM contained 1.94-fold higher Met than the EV control, whereas LM exhibited a 51.6% reduction relative to EV (Fig. 3).

**Figure 3.**
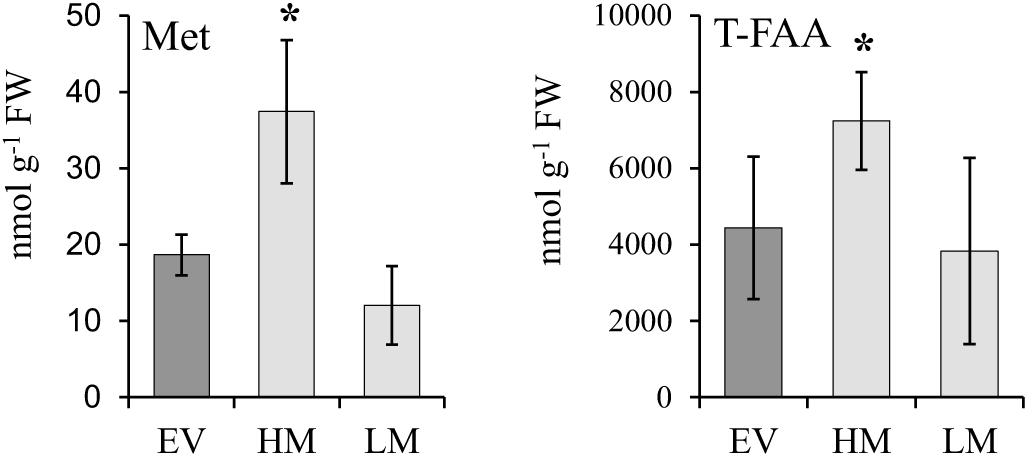
Met content and total free AAs (T-FAA) in leaves of SSE lines with high Met (HM), low Met (LM), and the empty vector (EV) control. Soluble Met and other AA levels were quantified in leaf tissue (fresh weight [FW]) using GC-MS and normalized to the ribitol internal standard. Data represents the mean ± SD of six biological replicates. Statistically significant differences compared to EV (p < 0.05, unpaired Student’s *t*-test) are indicated by asterisks.

Comprehensive primary metabolite profiling of HM, LM, and EV leaves was then conducted using GC-MS. Principal component analysis (PCA) revealed clear separation between HM and LM groups, indicating that Met levels strongly influence the overall primary metabolic profile in SSE leaves (Fig. S3A). Among the 15 detected AAs, 13 were elevated in HM compared to EV, whereas most were reduced in LM, apart from Ser and Gly (Table S1; Fig. S3B). Out of 39 quantified non-AA primary metabolites, 29 (74%) were reduced in LM, particularly sugars, compared to only 12 (30%) that were reduced in HM (Fig. S3C).

### High and low Met levels differentially affect gene expression

The pronounced metabolic differences between HM and LM plants suggest that altered Met levels may also influence gene expression. To test this, RNA sequencing (RNA-seq) was performed on the same leaf samples used for metabolic profiling (Table S2). Differential expression analysis relative to the EV control identified 1,022 differentially expressed genes (DEGs) in HM (representing 4.6% of detected genes), compared with only 107 DEGs in LM (0.47%) (Tables S2–S5; Fig. 4). Of these, 31 genes were commonly regulated in both SSE lines (Fig. 4A). Volcano plot analysis revealed a substantially larger number of both upregulated and downregulated genes in HM compared to LM, with the majority of DEGs in both cases being upregulated (Fig. 4B). To ensure robustness, only genes with normalized read counts greater than 20 were included in the analysis.

**Figure 4.**
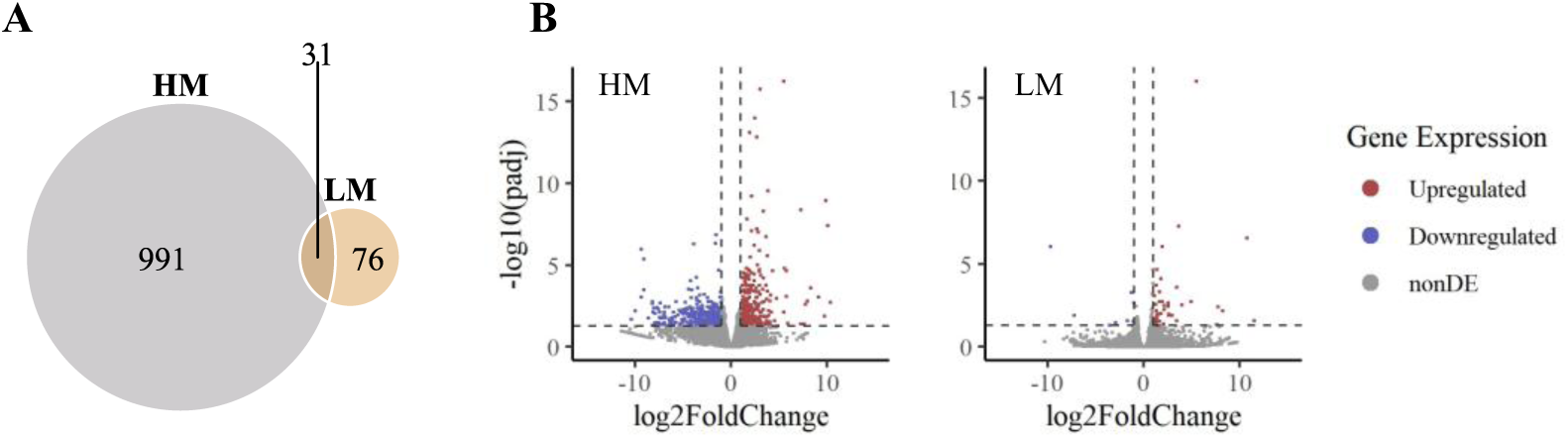
Summary of the RNA-sequencing analysis. (A) Venn diagram showing the overlap of DEG IDs between HM (gray) and LM (orange) SSE plants. (B) Volcano plots showing log₂ fold changes of DEGs in high-methionine (HM) versus low-methionine (LM) SSE lines relative to EV (padj < 0.05). Upregulated genes are shown in red; downregulated genes in blue.

### Transcriptome analysis reveals differences in Met pathway gene expression between HM and LM lines

To better understand the molecular mechanisms underlying the contrasting Met levels in HM and LM SSE lines, we examined the expression profiles of genes involved in Met metabolism using transcriptomic data. Our first focus was on *AtD-CGS* (Hacham et al., 2006). This form enables sustained Met synthesis under high Met conditions and is normally expressed at much lower levels than the full- length CGS in Arabidopsis (Hacham et al., 2006). Since *AtD-CGS* is highly expressed in SSE seeds (Cohen et al., 2014), we hypothesized that its expression in leaves could contribute to the observed differences in Met levels between HM and LM lines. Transcriptome analysis revealed that *AtD-CGS* transcript levels were 25-fold higher in HM than in LM, and 11.4-fold higher than in EV, whereas LM showed a 2.2-fold lower expression than EV (Fig. S4). The strong suppression in LM suggests the possibility of transcriptional or post-transcriptional silencing.

Although CGS is a central enzyme in Met biosynthesis, Met accumulation is also influenced by precursor availability and the activity of other enzymes in sulfur assimilation and carbon–amino acid pathways (Amir, 2010; Devi et al., 2023). Therefore, we extended our analysis to additional components of the Met metabolic network. In the sulfur assimilation pathway, most genes were downregulated in HM (Table S6; Fig. 5). For example, the sulfate transporter, *adenosine 5′-phosphosulfate reductases* (APR1– APR3), key enzymes in sulfate reduction (Koprivova and Kopriva, 2014), showed log₂ fold decreases of 1.8-3.6. In contrast, LM strongly upregulated these genes, including APS2, which catalyzes the conversion of APS to sulfite. Despite the overall suppression of sulfate assimilation genes, HM showed increased expression of *serine acetyltransferase* (SERAT1;1) and two isoforms of *O*-acetylserine(thiol)- lyase (OASTL A1 and OASTL B) (Koprivova and Kopriva, 2014), suggesting a compensatory boost in Cys biosynthesis to supply substrate for Met synthesis, likely driven by high AtD-CGS activity. In the Asp-family pathway, bifunctional *Asp kinase/homoserine dehydrogenases* (AK/HSDH I, II), which provide carbon–amino skeletons for Met biosynthesis (Wang et al., 2018), were downregulated in HM. *Met synthase 2* (MS2), which converts homocysteine to Met, was also suppressed (Table S6; Fig. 5).

**Figure 5.**
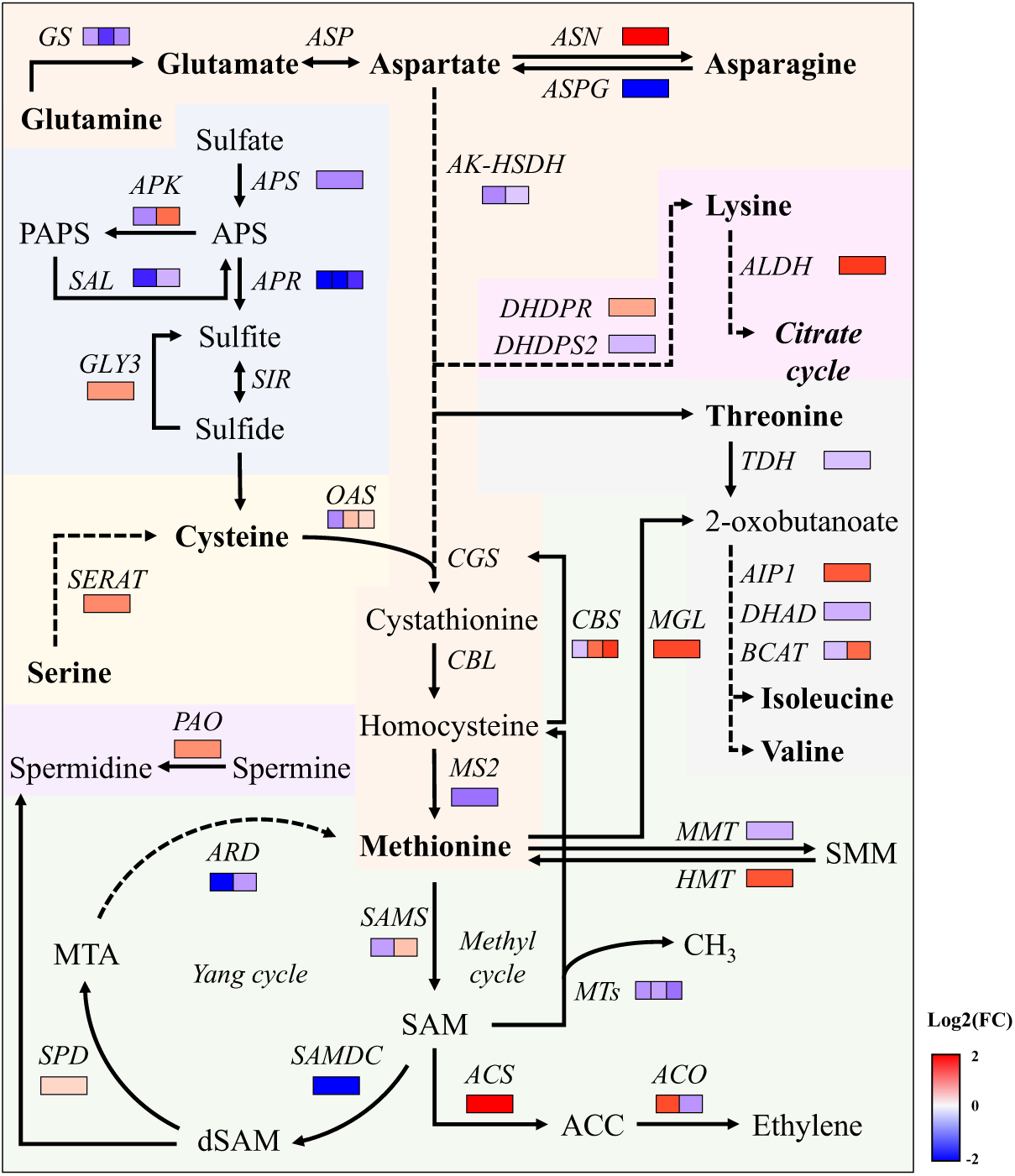
Differential expressions of genes involved Met metabolism in high-methionine (HM) versus low-methionine (LM) SSE lines. The diagram highlights key enzymes and metabolites involved in the metabolism of Met, including biosynthesis, degradation, and associated metabolic pathways. Only selected enzymes and intermediates are shown. Dashed arrows represent multiple enzymatic steps. Expression patterns are based on data presented in Table S11. Abbreviations: GS, glutamine synthase; ASN, asparagine synthase; ASPG, asparaginase; AK, aspartate kinase; DHPS, dihydrodipicolinate synthase; HSDH, homoserine dehydrogenase; CGS, cystathionine γ-synthase; CBL, cystathionine β- lyase; CBS, cystathionine β-synthase; MS, methionine synthase; SAM, S-adenosylmethionine; SAMS, SAM synthase; SAMDC, SAM decarboxylase; MTA, methylthioribose; SPD, spermidine synthase; ARD, acireductone dioxygenase; SAL, 3’(2’),5’-bisphosphate nucleotidase; MTs, methyltransferases; SMM, S-methylmethionine; MMT, Met S-methyltransferase; HMT, homocysteine S-methyltransferase; ACC, 1-aminocyclopropane-1-carboxylic acid; ACS, ACC synthase; EFE, ethylene-forming enzyme; MGL, methionine γ-lyase; TDH, threonine dehydratase; AIP, acute intermittent porphyria; DHAD, dihydroxyacid dehydratase; BCAT, branched-chain amino acid aminotransferase; APS, adenosine 5′- phosphosulfate; APR, APS reductase; APK, APS kinase; PAPS, 3′-phosphoadenosine-5′-phosphosulfate; SiR, sulfite reductase; GLY3, glycine; SERAT, serine acetyltransferase; OASTL, O-acetylserine (thiol) lyase; DHDPS, dihydrodipicolinate synthase; DHDPR, dihydrodipicolinate reductase; ALDH, aldehyde dehydrogenase.

Interestingly, HM leaves also showed signs of enhanced Met degradation. Met γ-lyase (MGL), which degrades Met to methanethiol, ammonia, and α-ketobutyrate (a precursor for isoleucine biosynthesis) (Goyer et al., 2007), was upregulated. Two key ethylene biosynthetic genes, *1- aminocyclopropane-1-carboxylic acid synthase* (ACS) and *ethylene-forming enzyme* (EFE), were also upregulated, along with 15 ethylene-responsive genes (Table S7, Fig. 5), suggesting increased ethylene production. In addition, *spermidine synthase* (SPDS2), a key enzyme in polyamine biosynthesis and the Yang cycle (Sauter et al., 2013), was upregulated in HM, supporting activation of SAM-utilizing pathways to reduce excess Met. However, most genes in the Yang cycle and methylation cycle were expressed at lower levels in HM than in LM, indicating that HM may limit Met recycling and resynthesis. Finally, *cystathionine β-synthase* (CBS) was predominantly upregulated in HM. Although its role in plants is not fully understood, in mammals, it participates in the Met catabolic pathway, diverting homocysteine toward Cys and promoting glutathione biosynthesis, potentially aiding in stress adaptation (Stipanuk and Ueki, 2011).

Aliphatic glucosinolates that are derived from Met are considered one of the main catabolites of Met. In Arabidopsis, up to 30% of the total sulfur pool is bound in glucosinolates (Falk et al., 2007). Met content is significantly increased in some aliphatic glucosinolate-deficient mutants (Chen et al., 2012; Shin et al., 2023). Determined the expression of genes in glucosinolate biosynthesis, showing that IMS2, the first enzyme committed to Met chain elongation, which facilitates an increased flux of Met into the glucosinolate pathway, and CYP79F1/2, which is involved in the formation of long-chain glucosinolates (e.g., 7MTH and 8MTO), were upregulated in both HM and LM (Table S9, Fig. S5). While in the HM line BCAT3, which channels the flux toward the short-chain aliphatic glucosinolate, was downregulated, in the LM line the expression of *IPMI1*, *CYP79F1*, *CYP83A1*, and *CYP83B1* were coordinated upregulation across multiple pathway branches (Table S9, Fig. S5). This suggests that the catabolism of Met to glucosinolates is one of the main factors that reduce Met in LM.

Taken together, these results suggest that HM respond to elevated Met levels by downregulating genes in both the Cys and Asp pathways that supply precursors for Met biosynthesis, while enhancing catabolic pathways to maintain Met homeostasis. In contrast, the reduced Met content in LM appears to relate to more catabolism to glcusinolates but might involve other regulatory mechanisms beyond transcriptional control of canonical Met biosynthetic genes.

### Met levels affect the expression of metabolism-related genes

Given the observed alterations in the primary metabolite profile (Fig. S3) and the differential expression of genes related to Met metabolism (Fig. 5), we further categorized DEGs associated with primary and secondary metabolism (Tables S8 and S9, respectively). Table S10A summarizes the number of DEGs grouped by metabolic clusters, as defined (Mukherjee et al., 2016). In both HM and LM lines, the most affected categories were AAs and sugar metabolism. HM and LM showed changes in 57 and 17 DEGs related to AA metabolism, and 51 and 17 DEGs related to sugar metabolism, respectively. In HM, most DEGs associated with AA metabolism were downregulated, with an average decrease of 2.3-fold (Table S10A). This reduction was observed across nearly all AA metabolic pathways (Table S10B). Interestingly, this transcriptional downregulation contrasts with the elevated AA levels detected in HM leaves. To address this apparent discrepancy, we examined DEGs associated with protein catabolism and found that 82% were upregulated (Table S10D), suggesting that the increased AA content may result primarily from enhanced protein degradation rather than *de novo* synthesis.

Similarly, many genes associated with sugar metabolism and polysaccharide catabolic processes were downregulated in HM (Tables S10A, C), including those involved in starch metabolism (Table S10D). The downregulation of starch metabolic genes suggests that fewer soluble sugars are incorporated into starch, potentially contributing to the higher glucose levels observed in HM.

In secondary metabolism, notable transcriptional changes were detected in the phenylpropanoid and terpenoid/carotenoid pathways. HM exhibited a greater number of both upregulated and downregulated DEGs compared to LM, with an overall trend toward downregulation in both lines (Table S10A).

### Differential expression of stress-related genes

To further investigate how altered Met levels influence gene regulation beyond primary metabolism, we performed Gene Ontology (GO) enrichment analysis of DEGs, focusing on the Biological Process category (Fig., Fig. S). In HM plants, upregulated genes were significantly enriched in stress-related pathways, including responses to desiccation, osmotic stress, insect attack, negative regulation of abscisic acid (ABA) signaling, and jasmonic acid (JA) biosynthesis (Fig.). Conversely, downregulated genes were associated with microtubule-based processes, ribosome biogenesis, gibberellin signaling, and other core cellular functions (Fig. S). In LM plants, upregulated DEGs were enriched for processes related to lysine and β-carotene metabolism, cell wall modification, tissue regeneration, seed morphogenesis, and response to water deprivation. Downregulated DEGs were enriched for translation, phospholipid transport, and nuclear pore organization (Fig. S).

The strong enrichment of ABA and JA signaling pathways in HM, both central regulators of plant stress responses, combined with elevated AA and sugar levels, which are commonly observed under abiotic stress (Salam et al., 2023), and with previous observations that SSE seeds accumulate higher transcripts of stress-related genes even under non-stress conditions (Cohen et al., 2014), prompted us to examine stress-related DEGs in more detail (Table S11). A total of 244 and 73 defense-related DEGs were identified in HM and LM, respectively, compared to EV (Table S11). Stress-related genes accounted for 23.9% of all DEGs in HM, nearly one-quarter of the transcriptomic changes, whereas LM contained only 8.85% stress-related DEGs. In HM, abiotic stress-related DEGs were 1.75-fold more abundant than those linked to biotic stress. In both SSE lines, upregulated stress-related DEGs outnumbered downregulated ones, with 1.77 times more upregulated genes in HM and 1.55-fold more in LM. Notably, HM exhibited 2.9 to 4-fold more upregulated and 2 to 5-fold more downregulated stress-related DEGs than LM (Table S11).

These transcriptomic changes may be attributed to enhanced expression of genes associated with four key stress hormones: ABA, JA, salicylic acid (SA), and ethylene (Table S7). Specifically, 108 DEGs related to ABA synthesis and response were identified in HM, of which 77 (71.3%) were upregulated. Similarly, among 91 JA-related genes, 67 (73%) were upregulated in HM. Of the 44 SA-related DEGs, 78.5% were upregulated. Likewise, 77 out of 98 ethylene-related DEGs (78.5%) showed increased expression (Table S7). These results strongly suggest that elevated Met levels are associated with broad transcriptional activation of stress hormone pathways and a wider stress-responsive gene network.

### DNA methylation analysis reveals hypermethylation in LM compared to HM

Our previous study indicated that DNA methylation levels are elevated in SSE leaves (Girija et al., 2023). The observation that Met content varied among individual plants within the same generation (Fig.) further suggests a potential link between Met levels and epigenetic regulation. This relationship is plausible given that SAM, derived from Met, serves as the universal methyl group donor for DNA methyltransferases (MTs) (Lashley et al., 2023).

To test this hypothesis, we performed whole-genome bisulfite sequencing (WGBS) on the same HM and LM leaf samples used for metabolomic and transcriptomic analyses, using the Illumina HiSeq 2500 platform (BGI Tech Solutions, Hong Kong). Sequencing achieved an average genome-wide mapping rate of 71% and bisulfite conversion efficiency >99% (Table S12A).

As expected, euchromatic regions displayed substantially lower methylation levels than heterochromatic regions, by approximately 4.4-, 10-, and 4.6-fold in the CG, CHG, and CHH sequence contexts, respectively (Table S12B), consistent with previous reports (Tan et al., 2016). When comparing SSE lines, LM consistently exhibited higher methylation levels than HM across all cytosine contexts (CG, CHG, CHH) and chromatin types (euchromatin and heterochromatin) (Table S12B).

To further explore these differences, we identified differentially methylated regions (DMRs) between HM and LM genomes. LM leaves contained more DMRs across all sequence contexts, with a total of 9,645 DMRs compared to 8,219 in HM, representing an ∼15% increase in methylation load (Fig. 6). The majority of DMRs occurred in the CHH context, followed by CG and CHG (Fig. 6). In both SSE lines, most CG-context DMRs were hypomethylated relative to EV, whereas CHG and CHH DMRs displayed a predominance of hypermethylation. Notably, LM showed a strong enrichment of hypermethylated CHG-DMRs (Fig. 6).

**Figure 6.**
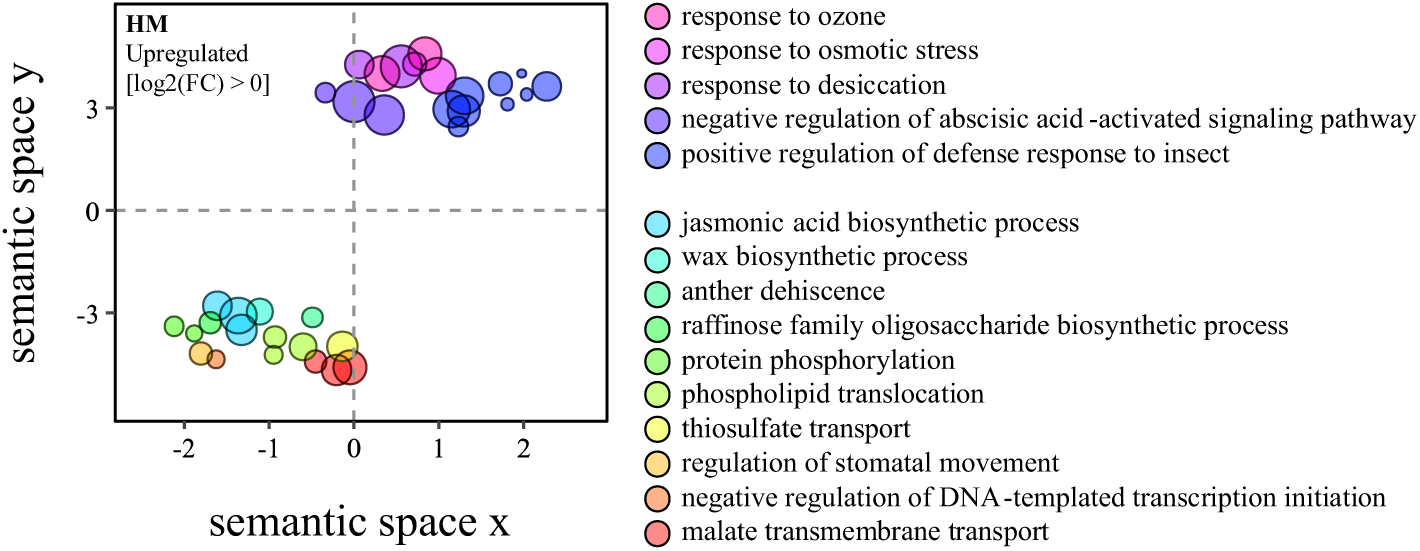
Multidimensional scaling scatter plot of GO enrichment analysis for Biological Process terms among upregulated DEGs in the high-methionine (HM) SSE line. Each circle represents an enriched GO term, with size proportional to statistical significance (larger circles indicate lower P-values). Colors group GO terms sharing similar parent GO categories.

To assess the genomic distribution of DMRs, Circos plots were generated for each methylation context (Fig. 6). CG-DMRs were broadly distributed throughout the genome in both lines, predominantly hypomethylated, and largely absent from centromeric and pericentromeric regions when compared to EV. CHG-DMRs, however, were highly enriched in centromeric and pericentromeric regions. In LM, most CHG-DMRs within centromeres were hypermethylated. In HM, CHG-DMRs extended beyond the centromeres into chromosome arms, showing a mixture of hyper- and hypomethylated regions. CHH-DMRs were dispersed across the genome in both lines, with higher densities in pericentromeric regions but clear depletion in centromeres. In both HM and LM, CHH-DMRs were predominantly hypermethylated. Overall, LM exhibited elevated non-CG methylation, particularly in CHG and CHH contexts within centromeric and pericentromeric regions. This enhanced methylation may contribute to the distinct metabolic and transcriptomic phenotypes observed in LM compared to HM.

### Association between DNA methylation and gene expression across genic features

Given the observed methylation changes in euchromatic regions (Fig. 6) and the substantial number of DEGs, we investigated the relationship between DNA methylation within gene bodies, associated regulatory elements, and normalized gene expression levels in HM and LM lines using a negative binomial regression model (Fig. S6). Across all cytosine contexts and genic features, correlation trends were generally similar between HM and LM; however, HM consistently exhibited stronger negative correlations for most genic features and methylation contexts, suggesting a more pronounced epigenetic influence on gene expression in the HM background.

**Figure 6a.**
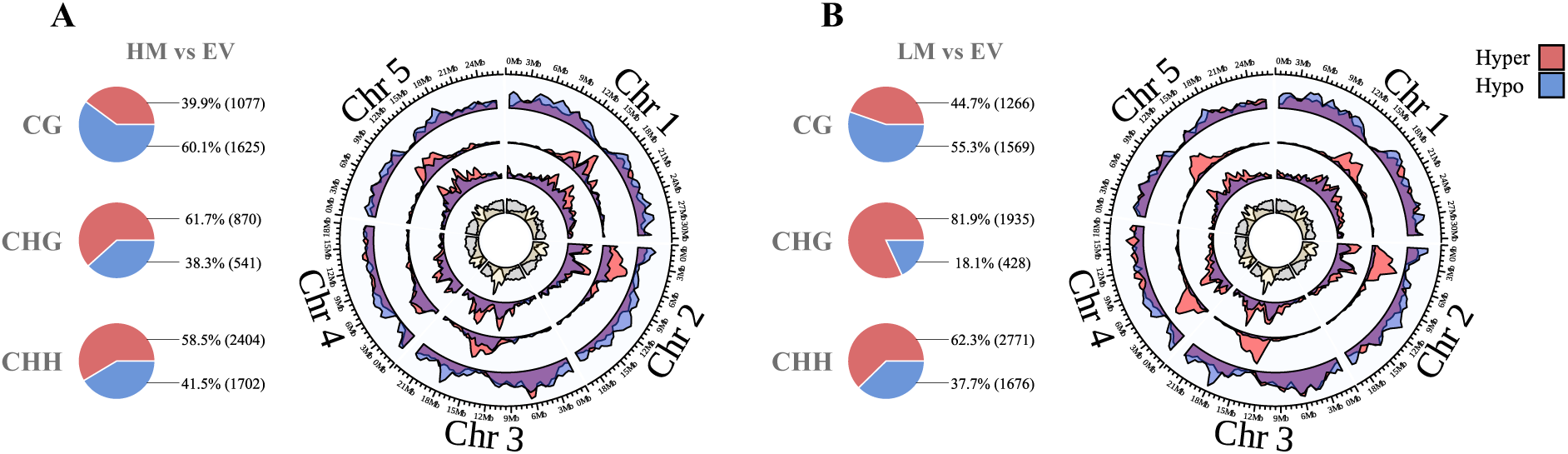
Differential methylation patterns in HM and LM compared to EV. (A, B) Pie charts show the proportions of hypermethylated (red) and hypomethylated (blue) differentially methylated regions (DMRs) in each cytosine context (CG, CHG, CHH) in HM (A) and LM (B) relative to EV control. The number of DMRs identified in each context is indicated in parentheses. Circos plots depict the genome-wide distribution of DMRs in CG, CHG, and CHH contexts (from outer to inner circles, respectively). Red indicates hypermethylated regions, blue denotes hypomethylated regions, and purple highlights areas where hyper- and hypomethylated DMRs overlap. The innermost circle shows the distribution of coding genes (gray) and the density of TEs (yellow), which mark pericentromeric heterochromatin regions.

Methylation levels were significantly associated with gene expression in all regression models. As a control, we examined TE-associated genes (TEGs, annotated in the TAIR10 database, www.arabidopsis.org), which are typically silenced by DNA methylation (Liu and Zhao, 2023). As expected, strong negative correlations between methylation and TEG expression were observed across all cytosine contexts (Fig. S7), validating the reliability of our model. Overall, the genome-wide methylation-expression relationship was comparable between HM and LM, suggesting that global methylation patterns alone cannot fully explain the transcriptomic differences between these lines. Instead, gene expression variation likely results from methylation changes in specific genomic regions.

We next quantified DMR distribution across coding sequences (CDSs), untranslated regions (UTRs), introns, and promoter regions. In both SSE lines, CG-context DMRs were most frequently located within CDSs, followed by introns, promoters, 3′ UTRs, and 5′ UTRs. HM exhibited slightly fewer CG-DMRs than LM (Fig. S8; Table S12), and both lines showed a general trend toward CG hypomethylation within genic regions, with HM displaying a slightly higher degree of hypomethylation in DEGs. In contrast, non-CG DMRs (CHG and CHH) were predominantly enriched in promoter regions and were mostly hypermethylated in both lines. LM displayed a higher number of promoter- hypermethylated non-CG DMRs than HM (Fig. S8). Since promoter hypermethylation, particularly in non-CG contexts, is generally associated with reduced gene expression in *Arabidopsis* (Zhang et al., 2018), this increased promoter methylation in LM may partially explain the lower magnitude of transcriptional changes observed in LM compared to HM. However, the differences between HM and LM in promoter methylation levels are modest, suggesting that additional regulatory mechanisms contribute to gene silencing in LM.

*AtD-CGS* transcript levels were 25-fold higher in HM than in LM (Fig. S3). To reveal if these differences are related to DNA methylation on the phaseolin promoter and/or AtD-CGS, we checked the differentially methylated positions (DMPs) across the phaseolin::AtD-CGS insertion in LM and HM lines (Fig. 7). Higher methylation was detected in the LM line compared to the HM line, with the LM having 26 DMPs, ∼2/3 higher than in HM that had 12 DMPs (Fig. 7B). Out of 12 DMPs, five were detected at the locus edges (promoter 5′ end and the 3×HA tag). However, the DMPs in LM were concentrated over the phaseolin promoter and the *AtD-CGS* gene body. By examining the contexts of the DNPs, a total of 38 DNPs were identified, of which 32 (89%) were CHH and 6 (11%) were CHG. The elevated methylation levels in the LM may indicate a partial epigenetic silencing of this region in the LM line.

**Figure 7.**
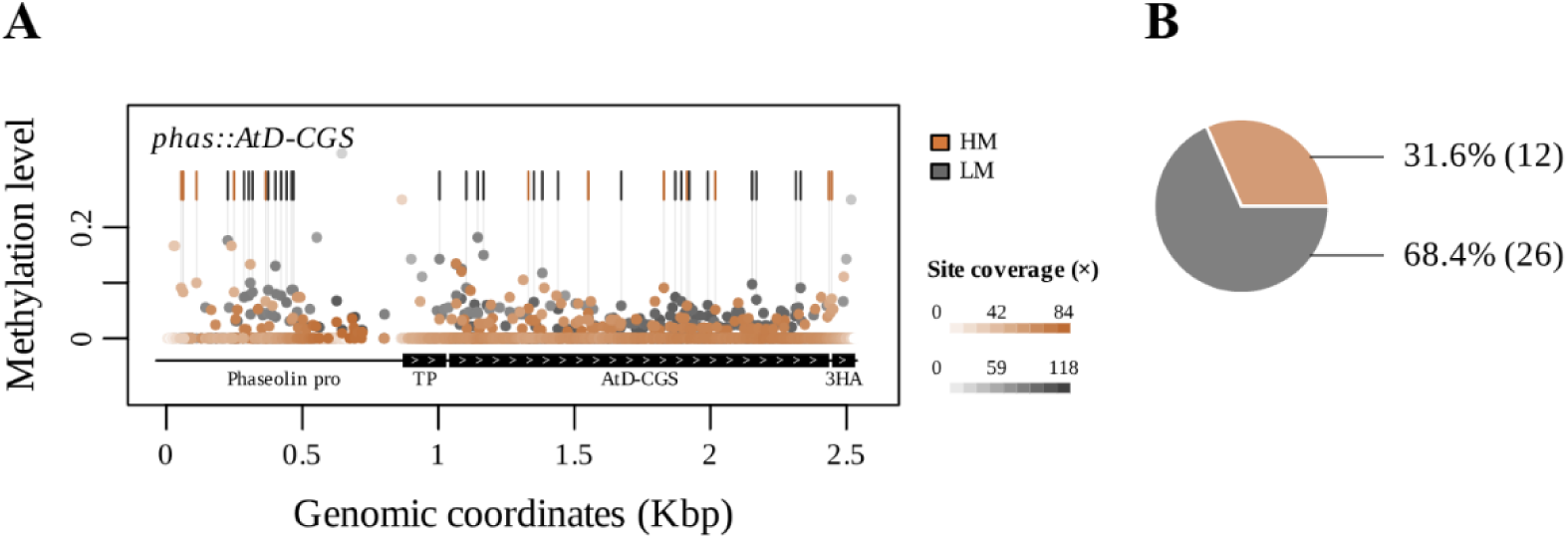
Differentially methylated positions (DMPs). (A) Methylation level over the phaseolin::AtD- CGS insertion. Each dot represents a single cytosine site in all contexts for HM (orange) and LM (gray). Darker dots indicate a larger read count. The vertical lines mark DMP sites that are significant. The Y- axis represents the methylation level, and the X-axis represents the genomic coordinates (Kb) over the phaseolin promoter, the rubisco small subunit transit peptide (TP), the AtD-CGS cDNA, and the 3HA- tag. (B) Pie chart summarizing DMP across all sequence contexts; the numbers show percentages from the total and counts.

### Altered expression of epigenetic machinery in HM and LM lines

The finding that LM plants exhibit higher methylation levels than HM plants, particularly in non-CG contexts, prompted us to compare the expression of genes associated with epigenetic regulation between the two lines (Table 1; Table S14). The analysis included genes encoding DNA methyltransferases (MTs), histone lysine MTs, components of the RNA-directed DNA methylation (RdDM) pathway, Royal family proteins, and chromatin remodeling factors (Tibben and Rothbart, 2024).

**Table 1.** Number of differentially expressed epigenetic regulator genes (*P* < 0.05, DESeq2) in high-methionine (HM) and low-methionine (LM) lines compared to the EV control. “Total” refers to the total number of genes identified for each gene group. Gene groups include DNA methyltransferases (MTs; EC 2.1.1.37), histone lysine MTs (Pontvianne et al., 2010), RNA-directed DNA methylation (RdDM) pathway components (Matzke & Mosher, 2014), Royal family proteins (Brasil et al., 2015), and other SAM-dependent methyltransferases.

| Group | Total Genes | HM / EV |  |  |  | LM / EV |  |  |  |
| --- | --- | --- | --- | --- | --- | --- | --- | --- | --- |
|  |  | Upregulated | Downregulated | Total Significant | Percentage | Upregulated | Downregulated | Total Significant | Percentage |
| DNA MTs | 8 | 0 | 3 | 3 | 37.50 | 0 | 1 | 1 | 12.50 |
| Histone-Lysine MTs | 46 | 0 | 6 | 6 | 13.04 | 1 | 3 | 4 | 8.70 |
| RdDM Pathway | 66 | 0 | 16 | 16 | 24.24 | 0 | 1 | 1 | 1.52 |
| Royal Family Proteins | 27 | 0 | 4 | 4 | 14.81 | 1 | 0 | 1 | 3.57 |
| Chromatin-Remodeling related | 92 | 0 | 15 | 15 | 16.30 | 0 | 4 | 4 | 4.35 |
| Other Methylation | 81 | 4 | 21 | 25 | 30.86 | 4 | 6 | 10 | 12.20 |

In HM, but not in LM, three DNA methylation-related genes were significantly downregulated: *maintenance DNA MT* (MET1), *cromomethylase 2* (CMT2), and CMT3. The expression levels of two DNA demethylases, *demeter-like 2* (DML2) and DML3, were also reduced (Table 1; Table S14). Among the six-histone lysine MTs examined, five showed reduced expression in HM. Similarly, 12 out of 13 DEGs involved in the RdDM pathway were downregulated in HM, along with three Royal family genes (Table S14). Furthermore, of the 25 DEGs encoding chromatin remodeling factors, 20 were downregulated in HM. These results indicate that in HM plants, where Met levels are elevated, there is a coordinated downregulation of key epigenetic regulators. This transcriptional suppression of DNA and histone methylation machinery may represent a compensatory mechanism to prevent excessive methylation activity, thereby maintaining epigenomic and transcriptomic homeostasis in the context of increased SAM availability.

## Discussion

This study reveals three key findings: (i) SSE plants expressing *AtD-CGS* under the seed-specific phaseolin promoter (Cohen et al., 2014) unexpectedly exhibited high *AtD-CGS* expression in leaves (Fig. 1, Fig. S1); (ii) Met levels in homozygous SSE plants from the third and fourth generations ranged widely from below empty vector (EV) controls in low-Met (LM) lines to approximately threefold higher in high-Met (HM) lines, with intermediate levels in the remaining lines; and (iii) HM and LM plants showed marked differences in their primary metabolomes, transcriptomes, and methylomes. These findings provide new insights into the molecular and epigenetic consequences of altered Met metabolism.

The unexpected *AtD-CGS* expression in SSE leaves is particularly noteworthy. The phaseolin promoter, which drives expression of a major seed storage protein in common bean (*Phaseolus vulgaris*), is generally considered embryo-specific and inactive in vegetative tissues (Chandrasekharan et al., 2003; Fait et al., 2011). However, our results strongly suggest that this promoter is sensitive to high Met and becomes active when the Met levels increase. Most probably, it occurs through chromatin remodeling. Indeed, a previous study has shown that its activity can be ectopically induced in Arabidopsis leaves via chromatin remodeling mediated by the *PvALF* (ABI3-like) transcription factor, especially under ABA treatment (Sundaram et al., 2013). In our SSE plants, transcriptomic analysis revealed no significant change in *PvALF* expression, suggesting that other regulatory mechanisms, possibly involving leaf- specific chromatin states or methylation patterns, account for the observed promoter activity.

The presence of both high and low Met levels among plants of the same generation and genetic background offered a unique opportunity to investigate the broader impacts of Met variation. The striking differences in T3 and T4 plants, nearly half of the progeny exhibiting Met levels lower than EV, and the remainder spanning a broad range, were unexpected. While homozygosity minimizes genetic variation, it does not preclude phenotypic variability arising from growth conditions and epigenetic regulation. Because all plants were grown under identical, controlled conditions, we attribute this variation primarily to epigenetic processes. The reduced Met content in LM plants likely reflects stochastic or progressive silencing of the transgene, potentially mediated by DNA methylation and/or histone modifications (Zhang et al., 2018). The observation that the phaseolin promoter and *AtD-CGS* gene in LM show higher changes in DNA methylation than HM, supports this assumption. Such epigenetic regulation is consistent with the metabolic role of SAM, the primary methyl donor and a direct catabolic product of Met. Elevated SAM availability can influence DNA and histone methylation, as well as other methylation-dependent processes, thereby linking Met metabolism to genome regulation (Roje, 2006). This potential feedback between Met levels and methylation provides a mechanistic framework for interpreting the distinct metabolic, transcriptomic, and methylomic profiles observed in HM versus LM lines.

### SSE lines with elevated Met content (HM) exhibit stress-associated shifts in both metabolism and gene expression

To characterize the physiological consequences of altered Met content, we profiled the primary metabolome, transcriptome, and methylome of HM and LM SSE lines. HM leaves exhibited elevated levels of both AAs and soluble sugars. Similar trends have been reported in seeds of *Arabidopsis*, soybean, and tobacco with enhanced Met content (Matityahu et al., 2013; Song et al., 2013; Cohen et al., 2014). Although the mechanisms underlying these increases remain unclear, both AAs and sugars are well-known to accumulate during abiotic stress (Salam et al., 2023). Consistent with this, HM leaves contained a substantially greater number of differentially expressed stress-related genes compared with both the EV control and LM lines (Fig. 4; Table S11), a pattern previously observed in SSE seeds with elevated Met (Cohen et al., 2014).

Specific sugars and AAs that accumulate under stress contribute to osmotic adjustment, reactive oxygen species scavenging, and serve as nitrogen storage or transport forms (Hildebrandt et al., 2015; Salam et al., 2023) (Fang et al., 2019). For example, biosynthetic pathways for Ser, Arg, Gln, and Ala are activated under osmotic and salt stress to provide precursors for rapid post-stress recovery (Batista-Silva et al., 2019). In parallel, stress conditions induce biosynthesis of AAs such as Pro, Gln, and Asn, whereas others (e.g., branched-chain AAs, Met, and Cys) are catabolized to support energy production (Hildebrandt, 2018). This coordinated regulation optimizes nitrogen and energy use during stress.

In HM leaves, most genes involved in AA biosynthesis were downregulated, while genes associated with protein catabolism were upregulated (Tables S8–S10), suggesting that the observed AA accumulation likely results from enhanced protein degradation. Additionally, the stress-associated reprogramming observed in HM, characterized by increased expression of ABA-, JA-, SA-, and ethylene-related genes, may further shift carbon/nitrogen metabolism toward osmolyte and defense compound synthesis (Sahu and Giri, 2025), thereby contributing to AA accumulation.

AAs also serve as precursors for major classes of secondary metabolites, phenylpropanoids, terpenoids, and alkaloids, that underpin plant defense against herbivores, insects, and pathogens, and contribute to tolerance of abiotic stress (Fang et al., 2019; Salam et al., 2023). In HM leaves, the majority of DEGs associated with terpenoid pathways were upregulated (65.4% relative to EV), whereas DEGs in phenylpropanoid/polyphenol pathways were predominantly downregulated (83% relative to EV) (Table S9). Stress-associated phytohormones, including ABA, JA, SA, and ethylene, are biosynthetically linked to or interact with these secondary metabolic pathways (Fang et al., 2019). Consistent with this, HM plants displayed elevated expression of most DEGs involved in the biosynthesis and signaling of these hormones (71–78% upregulated relative to EV), which likely contributes to the overall enrichment of stress-related transcripts (Fig. 4; Table S6).

Two Met-related processes directly connect to stress responses. First, ethylene biosynthesis frequently increases under stress, and Met serves as its precursor. Second, Met γ-lyase (MGL), a stress-inducible enzyme, degrades Met to produce methanethiol and Ile, metabolites implicated in stress adaptation (Goyer et al., 2007; Hacham et al., 2023). Although our data reveals a strong association between elevated Met levels and activation of stress-related transcriptional programs, the precise molecular mechanisms linking Met metabolism to these stress responses require further investigation.

### Met metabolism is differentially regulated in HM and LM lines

Given the low Met content in LM, we initially hypothesized that these plants would exhibit reduced expression of Met biosynthetic genes, with the opposite trend in HM. However, transcriptome profiling did not fully support this straightforward model. In HM, *AtD-CGS* expression was ∼25-fold higher than in LM, consistent with elevated Met levels; yet several key genes involved in *de novo* Met biosynthesis, spanning the Asp family pathway, sulfur assimilation, and Met synthesis (e.g., *APR*, *AK/HSDH*, *MS2*), were downregulated. This pattern suggests feedback regulation aimed at maintaining Met homeostasis (Fig. 4). Conversely, genes associated with Met catabolism and related pathways (e.g., *MGL*, *ACS*, *CBS*) and the genes related to glucosinolates were upregulated, indicating active buffering of excess Met (Hacham et al., 2002; Hacham et al., 2006; Goyer et al., 2007). This metabolic rebalancing likely contributes to the broader transcriptional reprogramming observed in HM, including activation of stress- response pathways and ethylene signaling components.

These results reinforce earlier findings that *AtCGS* overexpression prompts plants to dissipate surplus Met-derived carbon, methyl, and sulfur, for example, via dimethyl sulfide emission and enhanced ethylene production (Hacham et al., 2002). Elevated Met has also been linked to increased levels of catabolic products such as Ile and polyamines, as well as to the induction of glucosinolate biosynthetic genes (Cohen et al., 2014; Hacham et al., 2023). In addition, *S*-methylMet (SMM), a storage and transport form of Met, tends to accumulate under high-Met conditions (Boerjan et al., 1994). Together, these observations suggest that HM plants actively manage Met excess by channeling it into ethylene, polyamine, glucosinolate, and SMM pathways, and enhancing *MGL*-mediated catabolism. Notably, the reduced expression of genes in the Yang (Met salvage) cycle in HM suggests that decreased recycling of methylthioadenosine back to Met may also contribute to buffering excess Met.

The basis for the reduced Met content in LM, lower even than in EV control, remains unclear. It may reflect variation in upstream regulators of Met homeostasis, potentially involving master regulators of sulfur assimilation and the Asp family pathway. Identifying such regulators is important, as their manipulation can substantially increase Met accumulation. Despite recent advances in elucidating the regulatory network of Met metabolism (Devi et al., 2023), additional, as-yet-uncharacterized mechanisms appear to be involved. For example, a recent study in maize identified a post-translational regulator that deSUMOylates sulfite reductase (SiR) in the sulfur assimilation pathway, significantly increasing Met levels (Lu et al., 2025). Such findings indicate that further transcriptional and post- translational regulators of Met accumulation remain to be discovered.

### Met content affects DNA methylation levels in SSE plants

Given the extensive metabolic and transcriptional differences between HM and LM lines of identical genetic background, we next examined whether Met content influences DNA methylation. We performed WGBS to address this question. Epigenetic regulation and Met metabolism are closely interconnected in plants because SAM serves as the universal methyl donor for DNA and histone methylation. Reduced Met/SAM levels are associated with global DNA hypomethylation (Li et al., 2011; Groth et al., 2016; Meng et al., 2018; Yan et al., 2019), whereas engineered Met overaccumulation increases DNA methylation (González and Vera, 2019; Girija et al., 2023). This relationship is further supported by our recent work on the *mto1* mutant, which has elevated Met levels and displays DNA hypermethylation, particularly in non-CG contexts enriched in transposable elements (TEs), predominantly retrotransposon families (Yerushalmy et al. 2025).

WGBS revealed that, relative to HM, LM lines exhibit higher global DNA methylation, particularly in CHG and CHH contexts, with enrichment in centromeric and pericentromeric regions (Fig. 6). LM also displayed higher methylation levels and a greater number of DMRs in promoters (Fig. S7). In Arabidopsis, promoter hypermethylation is generally associated with reduced gene expression (Niederhuth et al., 2016; Zhang et al., 2018). However, this was not the prevailing pattern here: in both HM and LM, higher non-CG methylation in promoters was positively correlated with the increased expression of most genes (Fig. S6). Furthermore, the LM transcriptome was highly like that of EV, unlike HM (Table S2). Only a small subset of DEGs overlapped with DMRs in genic features in LM (Table S13B), suggesting that the modest transcriptional changes in LM are not directly explained by promoter methylation patterns. These results are consistent with previous reports (Bewick et al., 2017) that context- and region-specific methylation changes, rather than global methylation levels, are most relevant for gene regulation.

The most striking methylation differences between HM and LM were in CHH-context DMRs located in promoters (Table S13). In Arabidopsis, CHH methylation differs fundamentally from CG and CHG methylation, as it is established mainly *de novo* by DOMAINS REARRANGED METHYLTRANSFERASE 2 (DRM2) via the RdDM pathway and, in heterochromatin, maintained by CMT2 independently of RdDM (Law and Jacobsen, 2010) CHH methylation is asymmetric and must be re-established during each cell cycle, unlike CG and CHG methylation that are maintained through replication. Promoter-localized CHH methylation can recruit chromatin remodelers (Gallego-Bartolomé, 2020), potentially leading to transcriptional silencing via altered chromatin structure. Such effects could target *AtD-CGS* or other genes in Met biosynthesis, contributing to the reduced Met content and gene expression in LM.

In HM, a much larger proportion of DEGs overlapped with DMRs than in LM (Table S2), raising the question of whether DNA methylation plays a major role in the limited transcriptional changes in LM. The unexpected global hypermethylation in LM, despite low Met levels, suggests regulation beyond simple SAM pool size, possibly involving altered activity of DNA/histone methyltransferases and demethylases, changes in RdDM dynamics, or broader chromatin remodeling. Notably, we detected no methylation differences in the phaseolin promoter controlling *AtD-CGS* expression between HM and LM, consistent with previous suggestions that chromatin remodeling, rather than DNA methylation, regulates this promoter, as previously shown (Sundaram et al., 2013). This hypothesis could be tested using chromatin immunoprecipitation followed by sequencing (ChIP-seq) to profile histone modifications in SSE plants.

Interestingly, despite the higher DNA methylation in LM, relatively few epigenetics-related genes were differentially expressed compared to HM (Table 1). In contrast, HM exhibited widespread downregulation of epigenetics-related DEGs, including multiple DNA MTs and histone lysine MTs, consistent with a compensatory response to elevated SAM. This pattern supports the idea of a feedback loop linking central metabolism and chromatin state (Zhang et al., 2018), whereby HM plants actively downregulate methylation machinery to limit excessive methylation, while LM retains an epigenetic profile more similar to EV.

### A regulatory network integrating Met metabolism, stress responses, and epigenetic modulation

How elevated Met leads to the observed metabolic and stress-related phenotypes, and whether these effects are mechanistically linked to epigenetic regulation, remains to be fully elucidated. Such connections may involve as-yet-unidentified or unexpected regulatory nodes. One example of a plausible link is provided by a previously described network connecting JA, transcription factors, and Met metabolism. JA, a wound/herbivory signal who’s biosynthetic and signaling genes are upregulated in HM (Fig. 5), induces *MYB28*, which in turn activates core Met biosynthetic genes, including *CGS*, cystathionine β-lyase (*CBL*), and Met synthase (*AtMS2*), three key enzymes of the unique Met biosynthetic pathway (Hirai et al., 2007). *AtMS2* participates in recycling *S*-adenosylhomocysteine (SAH) to Met. Because SAH is a product of SAM-dependent MT reactions and a potent inhibitor of many MTs, including DNA MTs, JA–MYB28–Met crosstalk has the potential to influence chromatin methylation (Ouyang et al., 2020). In addition, MYB28 promotes aliphatic glucosinolate biosynthesis, a Met-derived defense pathway, and elevated Met may partly offset the diversion of carbon and sulfur into glucosinolates (Hirai et al., 2007).

In HM leaves, both *MYB28* and *LONG HYPOCOTYL 5* (*HY5*), a positive regulator of *MYB28* (Choi et al., 2024), were downregulated (Table S3). *HY5* is known to regulate *SULTR1;2* as well as *APR* and *ATPS1–ATPS3* in the sulfur-assimilation pathway (Huseby et al., 2013). The reduced expression of these genes in HM (Fig. 4) is consistent with attenuated sulfur uptake and assimilation under high-Met conditions. *HY5* also interacts with HISTONE DEACETYLASE 9 (*HDA9*) to modulate chromatin states at *HY5* target loci. *HDA9* acts on H3K9ac and H3K27ac, histone marks whose deacetylation is often associated with increased repressive histone methylation and transcriptional silencing (Yan et al., 2019). Downregulation of *MYB28*, *HY5*, and *HDA9* in HM may therefore represent a coordinated dampening of JA/HY5-dependent defense responses and associated chromatin programs. Future studies dissecting such complex interactions will be critical for understanding the yet-unresolved links between metabolic and transcriptional changes triggered by elevated Met.

In summary, while the effects of elevated Met levels on plant metabolism and gene expression have been extensively characterized in seeds (Matityahu et al., 2013; Song et al., 2013; Cohen et al., 2014; Zhang et al., 2023), their impact on leaf physiology remains comparatively underexplored. Investigating leaf responses is critical, as our previous work has demonstrated that leaves serve as a primary source of metabolites contributing to the enhanced nutritional quality of seeds (Girija et al., 2023). In the present study, we utilized genetically modified plants with either high-Met (HM) or low- Met (LM) content to examine the effects of elevated Met on the metabolome, transcriptome, and methylome. Our findings reveal that Met, beyond its canonical roles in translation initiation and methyl group donation, significantly alters plant phenotypes, with the most pronounced changes observed in HM lines. Notably, HM leaves exhibited increased levels of AAs and sugars, paralleling similar accumulations in seeds at the stage of SSE, thereby reinforcing the notion that leaves are central to metabolic provisioning during seed development (Girija et al., 2023).

The marked upregulation of *AtD-CGS* expression and accumulation of Met in HM appear to (i) downregulate genes in Met biosynthesis, (ii) upregulation of genes associated with Met catabolism, and (iii) reduce expression of genes involved in DNA methylation and chromatin remodeling. The enrichment of primary metabolites typically associated with stress responses, together with the elevated expression of stress-related genes in HM plants, suggests that high Met levels may induce a stress-like transcriptional state, even under non-stressful growth conditions. The molecular mechanisms underlying the connection between Met accumulation and activation of stress signaling pathways remain unclear and merit further investigation.

## Materials and Methods

### Plant genotypes and growth conditions

SSE and EV Arabidopsis plants were grown on 0.5× Murashige and Skoog (MS) medium (DuShefa, Haarlem, The Netherlands) supplemented with 1% (w/v) sucrose under a 16-h light/8-h dark photoperiod in a growth chamber at 22 ± 2 °C. After the initial growth phase, plants were transferred to soil and maintained under identical light and temperature conditions, as previously described by Cohen et al. (2014).

### Metabolite detection

Leaves from 21-day-old Arabidopsis plants were harvested, immediately frozen in liquid nitrogen, and lyophilized. The dried tissue was ground to a fine powder under liquid nitrogen and stored at –80 °C until analysis. Primary metabolites were profiled by gas chromatography–mass spectrometry (GC–MS) as previously described (Cohen et al., 2014; Girija et al., 2023).

### Whole-genome bisulfite sequencing (WGBS)

Genomic DNA was extracted from leaves of two biological replicates, each of HM, LM, and three biological replicates of the EV plants using a DNeasy Plant Mini Kit (Qiagen). Global DNA methylation profiles at single-nucleotide resolution were generated using the Illumina HiSeq 2500 platform (BGI Tech Solutions, Hong Kong). DNA quality and integrity were verified on 1% agarose gels.

For library preparation, 1 µg of genomic DNA per sample was fragmented to an average size of 200–350 bp, blunt-ended, A-tailed, and ligated to methylated adaptors. Libraries underwent bisulfite conversion using the EZ DNA Methylation-Gold Kit (Zymo Research), followed by PCR amplification, quality control, circularization, DNA nanoball (DNB) formation, and sequencing on the DNBSEQ platform (DNBSEQ Technology).

Raw reads were processed using SOAPnuke (Chen et al., 2018) with the following filtering parameters: removal of adaptor sequences (≥ 25% match, ≤ 2 mismatches), reads < 150 bp, reads containing ≥ 0.1% ambiguous bases (N), reads with polyX stretches > 50 bp, or reads with ≥ 40% of bases having Phred scores < 20. After filtering, only high-quality reads (Phred+33 encoding) were retained for downstream analysis.

### RNA sequencing (RNA-seq)

Total RNA was extracted from Arabidopsis leaves using the Spectrum Plant Total RNA Kit (Sigma- Aldrich) according to the manufacturer’s protocol. RNA integrity was confirmed by agarose gel electrophoresis and NanoDrop spectrophotometry. RNA-seq libraries were prepared at the Crown Genomics Institute of the Nancy and Stephen Grand Israel National Center for Personalized Medicine, Weizmann Institute of Science, using the INCPM-mRNA-seq protocol. Poly(A) RNA was purified from 500 ng total RNA, fragmented, and reverse transcribed to double-stranded cDNA. After Agencourt Ampure XP bead cleanup (Beckman Coulter), end repair, A-tailing, adaptor ligation, and PCR amplification were performed. Libraries were quantified using Qubit (Thermo Fisher Scientific) and TapeStation (Agilent). Sequencing was carried out on an Illumina NovaSeq 6000 platform (S1 flow cell, 100 cycles), generating 40–50 million paired-end reads per sample.

## Data analysis

For methylome analysis, clean reads were aligned to the *A. thaliana* reference genome (TAIR10) using Bismark (Krueger and Andrews, 2011). Differentially methylated regions (DMRs) were identified using the DMRcaller R package (Catoni et al., 2018), applying a beta-regression test for biological replicates in 100-bp bins with minimum proportion differences of 0.4 (CG), 0.2 (CHG), and 0.1 (CHH) contexts, requiring ≥ 4 cytosines per bin and ≥ 4 reads per cytosine.

For RNA-seq, clean reads were mapped to the Arabidopsis genome using RSEM (Li and Dewey, 2011) to obtain normalized expression values. Differentially expressed genes (DEGs; adjusted *p* < 0.05) were identified with DESeq2 (Love et al., 2014). Negative binomial regression was used to model normalized gene expression as a function of average methylation levels across gene bodies and promoters, accounting for genotype. Gene Ontology (GO) enrichment was performed with the topGO package using the ‘weight01’ algorithm (Alexa and Rahnenführer, 2007) and visualized using REVIGO (Supek et al., 2011). In the case of the normalized expression of the transgenic *AtD-CGS* gene in HM, LM, and EV (Figure S4), we quantified only the first 300-bp (using salmon [Patro et al., 2017]) encoding the plastid transit peptide of pea (*Pisum sativum*) rbcS-3A, which are absent from the endogenous *AtCGS* gene. Data visualization and statistical analyses were conducted in R (R Core Team, 2018). Custom R scripts are available at https://github.com/Yo-yerush.

## Data availability

All high-throughput sequencing data have been deposited in the NCBI BioProject database under accession number PRJNA1206675. Additional supporting data are available from the corresponding authors upon request. The Arabidopsis reference genome (TAIR10) and annotation files were obtained from The Arabidopsis Information Resource (www.arabidopsis.org).

## Acknowledgement

This work was supported by a grant from the Israel Science Foundation (ISF grant no. 1857/20).

## Author contributions

YH, and RA: experimental design. YY, MD, NR and YH: conducted experiments. YY and YH: data analysis. RA: preparing the first draft. YY, YH, RA: improved the manuscript preparation. All the authors read, revised, and approved of the manuscript.

## Supplemental Data

**Table S1.** Primary metabolite levels in leaves of HM and LM SSE plants compared to EV

**Table S2.** Summary output of the RNA-sequencing analysis of HM and LM SSE plants compared to EV

**Table S3.** DEGs overlapping DMRs in HM versus EV **Table S4**. DEGs overlapping DMRs in LM versus EV **Table S5.** DEGs overlapping DMRs in HM versus LM

**Table S6.** DEGs involved in Met metabolism in HM or LM vs EV and HM/LM.

**Table S7.** DEGs related to stress hormone pathways in HM and LM **Table S8.** DEGs, involved in primary metabolism in HM and LM **Table S9.** DEGs, involved in secondary metabolism in HM and LM

**Table S10.** Summary of DEGs that are associated with different processes of metabolism

**Table S11.** Summary of DEGs that are associated with stress

**Table S12.** Mapping and quality statistics for paired-end WGBS analysis

**Table S13.** Number of DMRs in various methylation contexts and gene features overlapping DEGs

**Table S14.** DEGs involved in epigenetic regulation in HM and LM

## Supplemental Figures

**Figure S1.** Elevated *AtCGS* expression, Met content, and total free amino acids (T-FAA) in SSE

**Figure S2.** Immunoblot detection of AtD-CGS protein in transgenic SSE Arabidopsis seeds

**Figure S3.** Primary metabolite profiles of SSE high-Met (HM), low-Met (LM), and empty-vector (EV) lines

**Figure S4.** Normalized expression of the transgenic *AtD-CGS* gene in HM, LM, and EV lines

**Figure S5.** Biological process enrichment for differentially expressed genes (DEGs) in HM and LM lines **Figure S6.** Relationship between average DNA methylation levels and normalized gene expression **Figure S7.** Annotation of DMRs in genic features of HM and L, SSE lines

